# Estimating the contribution of folding stability to non-specific epistasis in protein evolution

**DOI:** 10.1101/122259

**Authors:** Pouria Dasmeh, Adrian W.R. Serohijos

## Abstract

The extent of non-additive interaction among mutations or epistasis reflects the ruggedness of the fitness landscape, the mapping of genotype to reproductive fitness. In protein evolution, there is strong support for the importance and prevalence of epistasis, but whether there is a universal mechanism behind epistasis remains unknown. It is also unclear which of the biophysical properties of proteins—folding stability, activity, binding affinity, and dynamics—have the strongest contribution to epistasis. Here, we determine the contribution of selection for folding stability to epistasis in protein evolution. By combining theoretical estimates of the rates of molecular evolution and protein folding thermodynamics, we show that simple selection for folding stability implies that at least ~30% to ~60% of among amino acid substitutions would have experienced epistasis. Additionally, our model predicts substantial epistasis at marginal stabilities therefore linking epistasis to the strength of selection. Estimating the contribution of governing factors in molecular evolution such as protein folding stability to epistasis will provide a better understanding of epistasis that could improve methods in molecular evolution.

## INTRODUCTION

Epistasis refers to the non-linear and non-additive interactions among mutations. In the presence of epistasis, the genetic background affects the selective advantage of a mutation and the order at which amino acids are substituted matters. The extent of epistasis reflects the ruggedness and topology of the fitness landscape, which is the multi-dimensional mapping of genomic sequence to molecular properties and to organismal fitness. Since epistasis reflects the ruggedness of the fitness landscape, a mechanistic understanding of its origin is important for reconstructing the evolutionary history of proteins (<u>Harms and Thornton 2013</u>). Epistasis is also crucial to the predictability of microbial evolution, especially pathogenic bacteria and viruses (<u>O'Dea *et al*. 2010</u>; <u>Draghi and Plotkin 2013</u>; <u>Serohijos and Shakhnovich 2014b</u>; <u>Echave *et al*. 2016</u>; <u>Bershtein *et al*. 2017</u>; <u>LÄssig *et al*. 2017</u>). Additionally, understanding the extent and mechanism for epistasis is crucial for inference of evolutionary past, such as phylogenetic methods and various statistical tests for adaptive evolution (<u>Weinreich *et al*. 2013</u>). A major shortcoming of most of the methods in molecular evolution is the assumption of additivity of mutational effects and independent evolution among sites within a gene or protein. Accounting for epistasis could enhance the accuracy and predictability of these tools (<u>Cordell 2002</u>).

There are numerous examples of epistasis in proteins (<u>Starr and Thornton 2016</u>) (<u>Miton and Tokuriki 2016</u>). In the evolution of cefotaxime resistance in *E. coli* ß-lactamase, mutations that enhanced cefotaxime hydrolysis are also destabilizing and therefore only beneficial in high stable backgrounds (<u>Bershtein *et al*. 2006</u>; <u>Weinreich *et al*. 2006</u>). Similar findings were observed in the evolution of the vertebrate glucocorticoid receptor (<u>Bridgham *et al*. 2009</u>) and the nucleoprotein in human influenza viruses (<u>Gong *et al*. 2013</u>). In the study of site-directed mutagenesis in hepatitis C virus NS3 protease variants, the same mutations introduced to different backgrounds resulted in a broad range of fitness effects, from nearly-neutral to almost lethal (<u>Parera and Martinez 2014</u>).

In general, non-commutativity of mutations can be of two types—*magnitude epistasis* and *sign epistasis*. In magnitude epistasis, the beneficial or deleterious nature of mutations remains unchanged but their magnitude is amplified or suppressed depending on genetic background. However, in sign epistasis, the beneficial/deleterious nature of mutations are interchanged, a beneficial mutations in one genetic background can become deleterious in another (<u>Weinreich *et al*. 2005</u>). By comparing stability effect of all single and double mutations of IgG-binding domain of protein G, Olson et al. reported pervasive sign epistasis among different combinations of mutations (<u>Olson *et al*. 2014</u>). Epistasis can also be classified as either positive or negative. *Positive epistasis* occurs when the combined effect of two mutations is higher the arithmetic sum of their individual effects, *negative epistasis* occurs when this sum is lower.

In proteins, it is helpful to distinguish between *specific epistasis* and *non-specific epistasis* (<u>Starr and Thornton 2016</u>) (Figure 1). Specific epistasis results from direct physical interaction of spatially close residues in 3D structure or indirect influence of residues on each other through long-range allosteric effects. Mutations occurring at spatially proximate sites will have non-additive contribution to the biophysical properties of proteins such as stability, activity, dynamics, or binding with partner proteins (<u>Studer *et al*. 2013</u>). If the biophysical properties determine organismal fitness, as recently shown in examples of viral and microbial evolution (<u>Bershtein *et al*. 2017</u>), the non-additivity at the level of proteins translates to non-additivity at the level of fitness. As a consequence, the rate and patterns of substitution in one site may be correlated with that of another spatially close site (<u>SÜel *et al*. 2003</u>; <u>Morcos *et al*. 2011</u>; <u>Marks *et al*. 2012</u>; <u>Pollock *et al*. 2012</u>; <u>Dickinson *et al*. 2013</u>).

**Figure 1.**
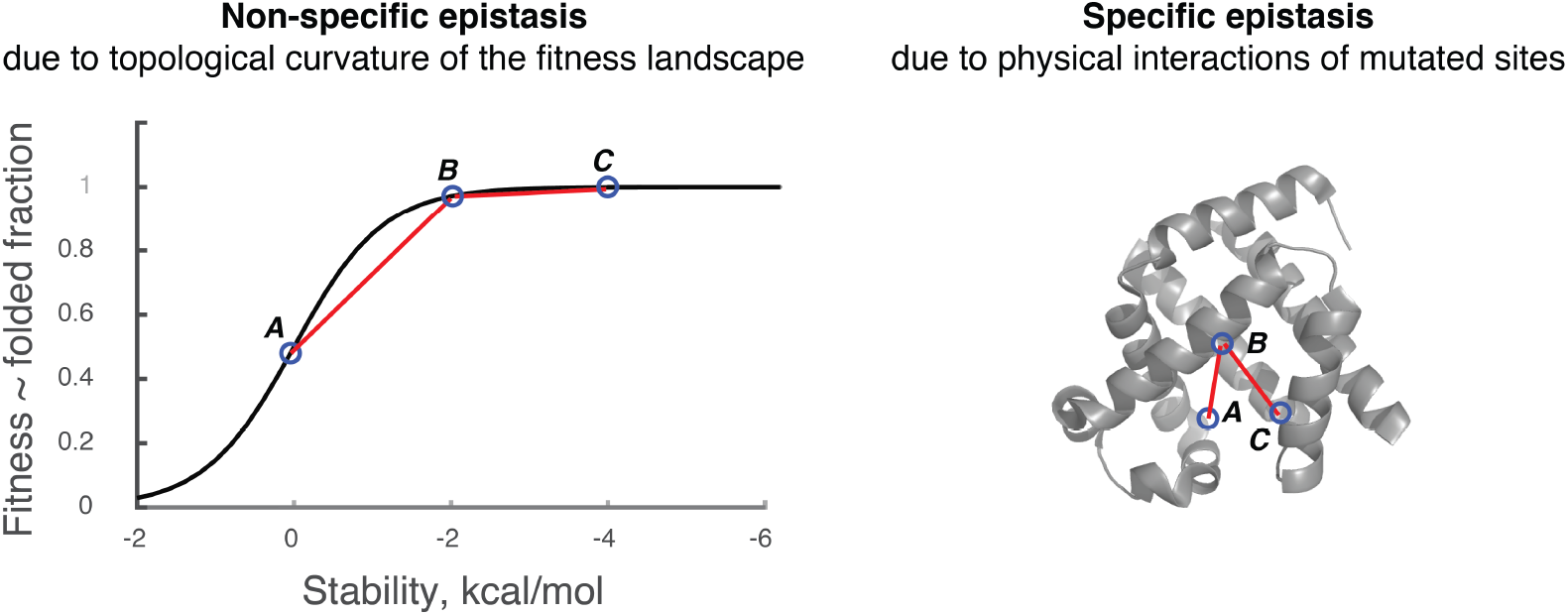
**Non-specific epistasis.** Non-specific epistasis is defined as non-linear relationship between mutational effects on the structure and protein property, e.g., protein stability and between protein properties and cellular fitness. In the figure, evolution is epistatic with respect to the order of A, B and C mutations even though they might be located far from each other on 3D structure.

Non-specific epistasis arises from the non-linear dependence of cellular/organismal fitness to biophysical properties such as folding stability (Figure 1). Even if biophysical traits are additive and non-epistatic, the non-linear mapping of the biophysical property to fitness introduces non-linear interactions among mutations at the fitness level. Since Darwinian selection acts at the level of organismal fitness, non-specific epistasis could affect the rate of evolution. The simplest mapping between fitness and protein properties exhibits a single peak, such as shown in Figure 1 for folding stability. This single-peak and plateau-like fitness landscape has also been shown for the relationship between fitness and intracellular abundance of a gene and between fitness and enzyme activity (<u>Flint *et al*. 1981</u>; <u>Dykhuizen *et al*. 1987</u>; <u>Bershtein *et al*. 2013b</u>; <u>Bershtein *et al*. 2013a</u>; <u>Bershtein *et al*. 2015b</u>; <u>Rodrigues *et al*. 2016</u>). In these simple landscapes, non-specific epistasis gives rise to the “law of diminishing returns”, i.e., the selective advantage of mutations decreases as the fitness of the organism increases, a well-known feature of many optimization processes and in adaptive trajectories in protein evolution (<u>Hartl *et al*. 1985</u>; <u>Chou *et al*. 2011</u>; <u>Miton and Tokuriki 2016</u>).

Several works have investigated the role of stability to protein epistasis (<u>Bershtein *et al*. 2006</u>; <u>Wylie and Shakhnovich 2011</u>; <u>Ashenberg *et al*. 2013</u>; <u>Gong *et al*. 2013</u>; <u>Serohijos and Shakhnovich 2014a</u>; <u>Shah *et al*. 2015</u>; <u>Bershtein *et al*. 2017</u>). Folding stability is a universal property of proteins that determine the fraction of proteins in the native state. Based on the simple assumption that proteins need to be folded for organisms to be viable, a quantitative fitness landscape (Figure 1) may be constructed from protein folding thermodynamics (protein folding fitness landscape or PFL). Folding stability directly affects the evolutionary rate of proteins (<u>DePristo *et al*. 2005</u>; <u>Bloom *et al*. 2006</u>; <u>PÁl *et al*. 2006</u>; <u>Serohijos *et al*. 2012</u>; <u>Serohijos and Shakhnovich 2014b</u>; <u>Echave *et al*. 2016</u>; <u>Bershtein *et al*. 2017</u>). Gain-of-function mutations are on average destabilizing (<u>Tokuriki *et al*. 2008</u>), thus stabilizing substitutions can act as permissive or compensatory mutations that increase the likelihood of fixation of functional mutants (<u>Weinreich *et al*. 2005</u>; <u>Soskine and Tawfik 2010</u>; <u>Gong *et al*. 2013</u>). Additionally, several observations in molecular evolution and genomics have been explained based on PFL (<u>Serohijos and Shakhnovich 2014b</u>). For example, folding stability has a direct role in the genomic observation that highly abundant proteins evolve slowly (<u>Serohijos *et al*. 2012</u>), a consistent finding across organisms from different kingdoms of life (<u>Drummond and Wilke 2008</u>).

Despite these works highlighting the role of stability in protein epistasis, several questions are unanswered. First, to *what extent does selection for folding stability contribute to protein epistasis?* Kondrashov and collaborators argues that epistasis is pervasive in protein evolution and estimated that up to ~90% of amino acid substitutions experienced epistasis (<u>Breen *et al*. 2012</u>). However, it is not established what factors could give rise to this prevalence of epistasis.

Here we hypothesize that this estimated fraction of epistasis is systematically influenced by selection for folding stability. To elucidate the contribution of PFS to epistasis, we build on the previous work of Breen *et al*. that estimated epistasis by comparing two relative evolutionary rates—the average dN/dS among protein orthologues and the average mutational usage (Figure 2A). Using a combination of forward evolutionary simulations and theoretical analysis, we show that the fraction of amino acid substitutions that experience epistasis due to folding stability is at least ~30%, and could reach up to ~60% for proteins evolving at low stability. We also find significant negative epistasis under selection for folding stability, in agreement with experimental observations. The magnitude of this epistatic interactions are increased at marginal folding stabilities. Since marginal stability is also the regime where purifying selection is strong, our results highlight the strong coupling between epistasis and the strength of selection. Altogether, our quantitative estimate of the contribution of selection for folding stability to epistasis, could lead to a more mechanistic understanding of this important evolutionary force.

**Figure 2.**
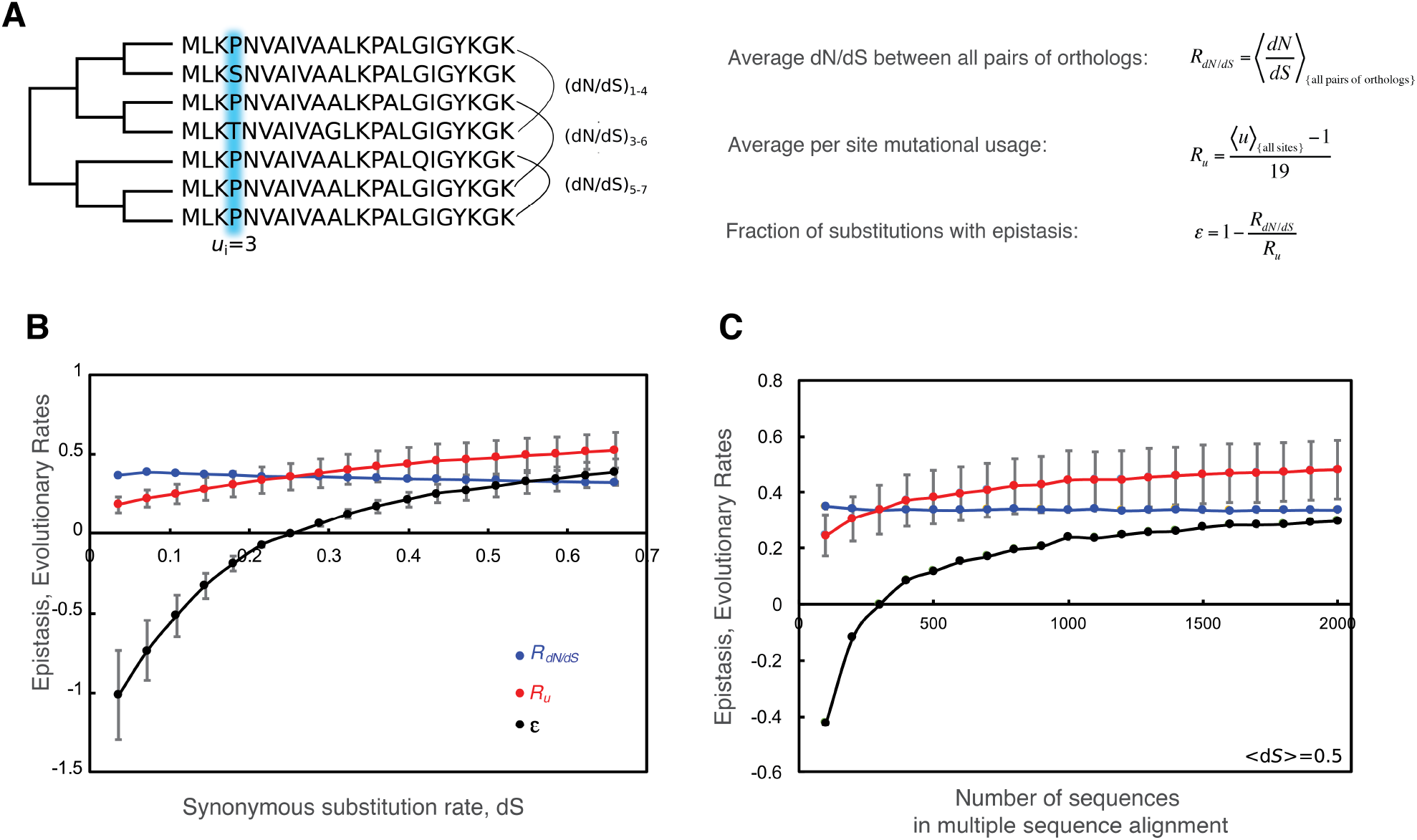
**Protein sequences evolved under selection for thermodynamic stability show on average 30% epistasis.** **(A)** Orthologous protein sequences are used to estimate epistatic evolution as the between rate of evolution over all possible backgrounds estimated from mutational usage, R_u_, and pairwise rate of evolution, *R*_dN/dS_. **(B)** *R*_dN/dS_ and Ru are plotted as a function of sequence divergenc. **(C)** Pairwise rate of evolution, *R*_dN/dS_ and the rate calculated from mutational usage, R_u_, are plotted for different number of sampled sequences corresponding to the MSA at <d*S*>=0.5 in panel B. In both B and C, epistasis is calculated as the percentage difference between the two rates, *R*_dN/dS_ and Ru.

## RESULTS

### Non-specific epistasis due to the protein folding fitness landscape

In the absence of epistasis, the substitution rates are independent of genetic background. Thus, to estimate epistasis, one approach is to compare the rates of substitution of mutations with and without background specificity. Kondrashov and coworkers (<u>Breen et al. 2012</u>) applied this method to estimate the extent of epistasis in protein evolution by comparing two rates calculated from a multiple sequence alignment (MSA) of orthologs—the average pairwise substitution rate *R*_dN/dS_ and the rate of mutational usage *R_u_*:

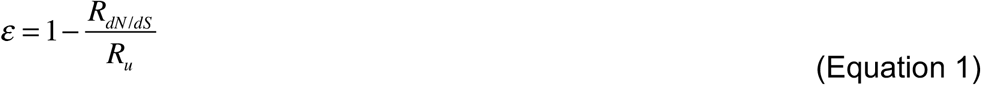

*R*_dN/dS_ is the average d*N*/d*S* (the ratio of non-synonymous substitution rate *dN* and synonymous substitution rate *dS*) for all pairs of orthologues in an MSA. *R*_dN/dS_ reflects background- and lineage-specificity of amino acid substitutions (Figure 2A). 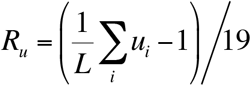 where *u* is the number of unique amino acids in each site in an MSA and is referred to as the mutational usage. *L* is the length of the protein. *R_u_* represents the ratio between observed accessible amino acid substitutions in a site, (*u*-1), and all possible amino acid substitution assuming no selection, that is, (20-1)=19. Because *R*_u_ is calculated per site, it is independent of background and lineage.

Selection is major determinant of substitution rates, thus any estimate of epistasis must normalize for the confounding role of selection. *R*_dN/dS_ reflects selection because it is simply the pairwise dN/dS between orthologs and *dN*/*dS* itself is a measure of the stringency of selection. *R_u_* also reflects selection. Without selection and if all mutations are neutral, all 20 amino acids are accessible in each site, thus *R_u_*=1. Altogether, *R_u_* represents the optimal evolutionary rate with selection but without epistasis, while *R*_dN/dS_ is the observed rate that fully reflects both selection and epistasis. Hence, Equation 1 includes the normalization for the confounding effect of selection.

From a set of diverse proteins, Kondrashov and co-workers (<u>Breen *et al*. 2012</u>) calculated *R*_dN/dS_ ~ 0.01-0.1 and *R*_u_ ~ 0.1-0.6, resulting in an estimate of epistasis to be *ε*~0.6-0.9. This finding implies that about ~60% to ~90% of amino acid substitutions in proteins experienced epistasis. Despite this inferred prevalence of epistasis in long-term protein evolution, a universal mechanism, if any, is lacking.

To determine how much epistasis can be explained by folding stability, we simulate protein sequence evolution under selection for stability, generate multiple sequence alignment, and apply Equation 1. Briefly, protein sequences are evolved using a Wright-Fisher sampling approach with the fitness function:

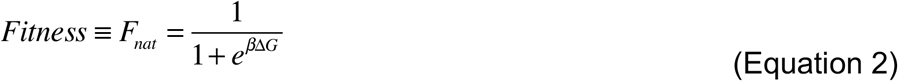

where Δ*G* is the folding free energy and *β* is the Boltzmann constant. Equation 2 represents the probability that a protein is in the native state, which is required for function. When a random non-synonymous mutation occurs, it changes the folding stability of the wildtype by ∆∆*G* = ∆*G_mut_* − ∆*G_WT_*, where Δ*G_mut_* is the new stability of the mutant. In this folding stability fitness landscape, the selection coefficient is

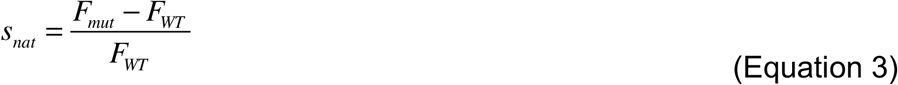

The functional form of *s_nat_* distinguishes between different background stabilities as widely discussed before (<u>Chen and Shakhnovich 2009</u>; <u>Goldstein 2011</u>; <u>Serohijos and Shakhnovich 2014b</u>). Assuming that the population is monoclonal, at each mutational attempt, the probability of fixation is defined by the Kimura formula,:

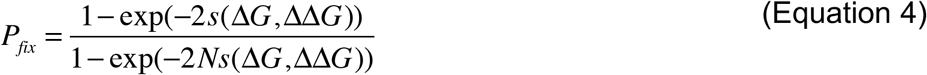

In our simulations, we assume an effective monoclonal population size of *N_e_*=10^4^. Our model system is *dihydrofolate reductase* (DHFR) taken from *Candida Albicans* with PDB ID=1AI9 (<u>Whitlow *et al*. 1997</u>). The PDB structure is used to estimate the effect random mutations on folding stability DD*G* using a physical force field (Methods).

First, we simulate 2000 independent trajectories of protein evolution all starting from the same initial ancestral sequence. This procedure mimics the divergence of orthologs from a common ancestor. We run the simulation for 10^7^ generations and extracted nucleotide sequences every 10^5^ generations (that is, every 10*N_e_*), thus creating a set of MSAs with different divergence times. Estimating epistasis on this set of MSAs controls for the dependence of both *R_u_* and *R_dN/dS_* on divergence time (Figure 2B). We use average pairwise synonymous substitution rate as a measure of divergence time and show the range of <*dS*> up to 0.5 which is the same range of <*dS*> for the protein set in (<u>Breen *et al*. 2012</u>) (Figure 2B and Table S1). *R_u_* increases almost three folds from 0.18 (~five unique amino acids per site) to 0.6 (~13 amino unique acid per site). *R_dN/dS_* is slightly higher at initial divergence time due to stochasticity in *dS* because of few fixed synonymous substitutions. Estimated epistasis based on Equation 1 indeed shows a strong dependence on divergence time (Figure 2B). For the set of proteins studied in (<u>Breen *et al*. 2012</u>) <*dS*>=0.4, which from our simulation using folding stability, the estimated epistasis is ~30%. Thus, while Kondrashov and co-workers estimated that ~60% to ~90% epistasis in protein evolution (<u>Breen *et al*. 2012</u>), we find that up to half of this estimated can be accounted for by simple selection for folding stability.

Additionally, in our simulations based on folding stability, the mutational usage is *u*~13, which means that each site in the MSA has an average of 13 (out of the possible 20) unique amino acids. For the set of proteins studied in (<u>Breen *et al*. 2012</u>), *u*~8. This difference is expected because real protein sequences are under selection for other biophysical properties (e.g., protein-protein interaction, binding, dynamics) and biological function beyond folding stability. Interestingly, the difference between u=20 (i.e., full mutational usage and zero selection), u=13 (under selection for thermodynamic stability) and u=8 (in real proteins) provides a rough estimate of the contribution of protein folding stability to selection relative to other properties. That is, the drop in amino acid usage per site in an MSA, (20-13)/(20-8) or ~60% is due to selection for folding stability.

To control for the potential bias of the number of sequences in the MSA to Equation 1, we down-sample the sequences and estimated epistasis (Figure 2C). We perform this down-sampling procedure on the most diverged MSA corresponding to <*dS*> = 0.5. *R*_u_ converges to ~0.65 (u~13) while *R*_dN/dS_ is ~0.30 at all number of sampled sequences. Therefore, epistasis converges to ~ 30% once enough sequences (>1000) are used in the MSA and the estimation of *R_u_* and *R_dN_*_/*dS*._

### Prevalence of negative epistasis under selection for thermodynamic stability

In the analysis above, we examine the role of epistasis on long-term protein evolution by analyzing amino acid substitutions across orthologues. Next, we analyze short-term epistasis by focusing on the pairwise interaction between two randomly arising mutations. Such an analysis is directly comparable to results from directed mutagenesis or comprehensive deep mutational scans. Specifically, we want to determine which type of pairwise epistasis, positive or negative, dominates under selection for stability. To do so, we pick two random mutations *A* and *B* that change the folding stability of wildtype Δ*G_WT_* by ΔΔ*G_A_* and ΔΔ*G_B_*, respectively. These values are drawn from the distribution of random effect of mutations on folding stability *p*(*∆∆G_fold_*) inferred from the collection of experimental measurements and comprehensive computational mutagenesis of globular proteins (<u>Tokuriki *et al*. 2007</u>). Using Equation 1, the corresponding fitness of these two mutants are *F_A_* and *F_B_*. Since we focus only on non-specific epistasis, their combined effect on stability is ΔΔ*G_A_* + ΔΔ*G_B_* with a corresponding fitness value *F_AB_* = *F*(ΔΔ*G_A_* + ΔΔ*G_B_*). We calculate the pairwise epistasis using the following equation:

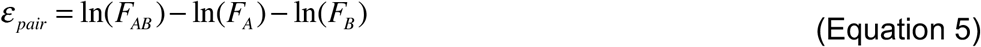

where *F_A_* and *F_B_* are fitness of each mutant. We show in Figure 3A-B the pairwise epistasis among random mutations as a function of background folding stability. Indeed, the most dominant form of epistasis is negative, in agreement from comprehensive mutational scans in IgG-binding domain of *Streptococcus* protein G (<u>Olson *et al*. 2014</u>), substrate binding domain of yeast Hsp90 (<u>Bank *et al*. 2015</u>) and green fluorescent protein from *Aequorea victoria* (<u>Sarkisyan et al. 2016</u>). This negative epistasis is largely due to pairs of mutations that are destabilizing (Figure 3C). We also plotted epistasis versus the ratio of ΔΔG values of pair mutations in Figure S1. Since destabilizing mutations bring the protein to the more curved part of the PFL, negative epistasis starts at high stability compared to positive epistasis (Figure 3B). Altogether, the PFL features greater curvature near low folding stability, epistasis is larger in the regime where selection is stronger.

**Figure 3.**
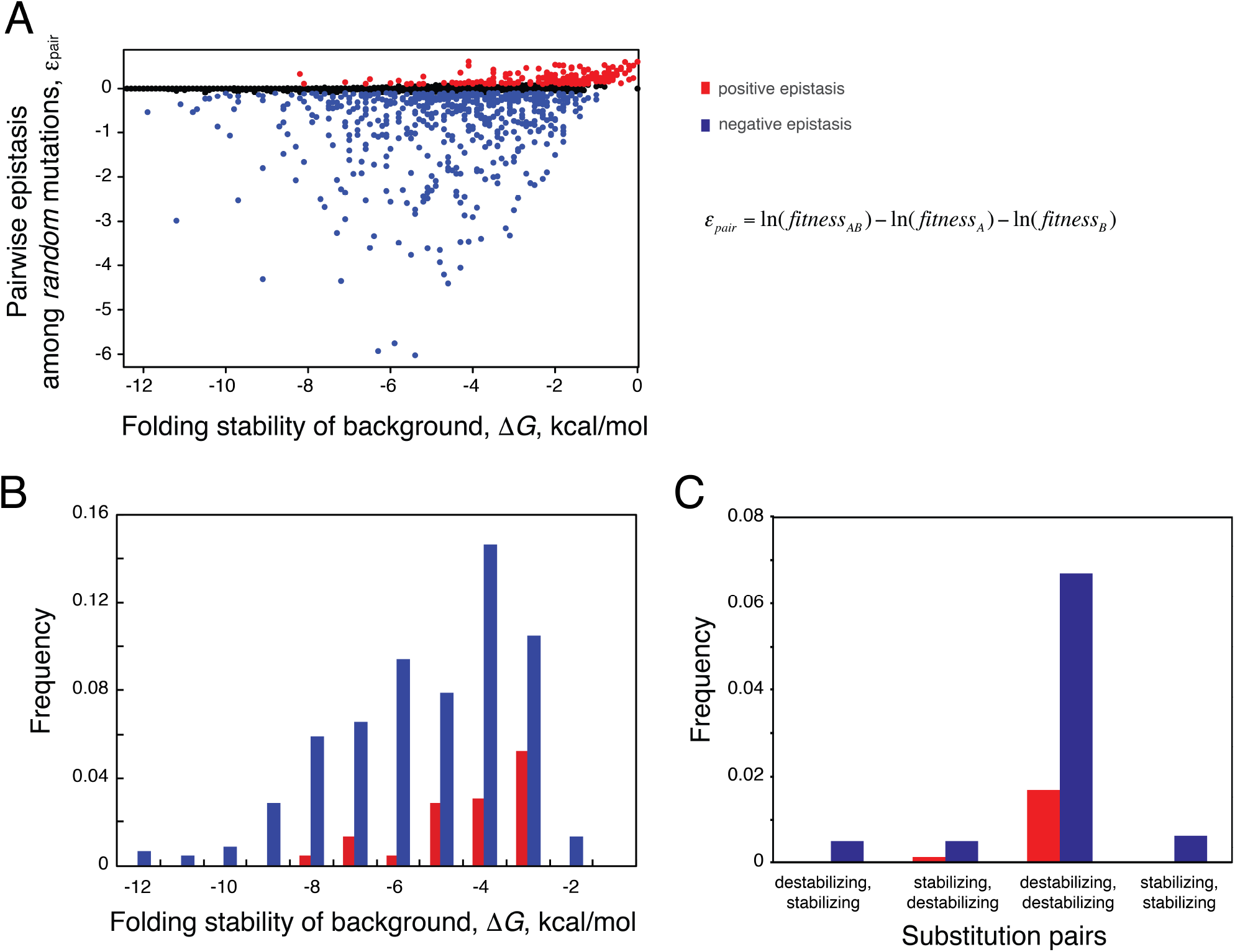
**Pervasive negative epistasis under selection for thermodynamic stability.** **(A)** Pair epistasis between two randomly chosen mutations *A* and *B* as a function of the folding stability of the background. Positive epistasis ε_pair_>0.1 are shown in red; Negative epistasis ε_pair_<-0.1 are shown in blue. **(B)** Frequency of positive and negative epistasis with respect to wildtype stability. **(C)** Frequency of positive and negative epistasis with respect to effect of mutations on stability.

**Figure 4.**
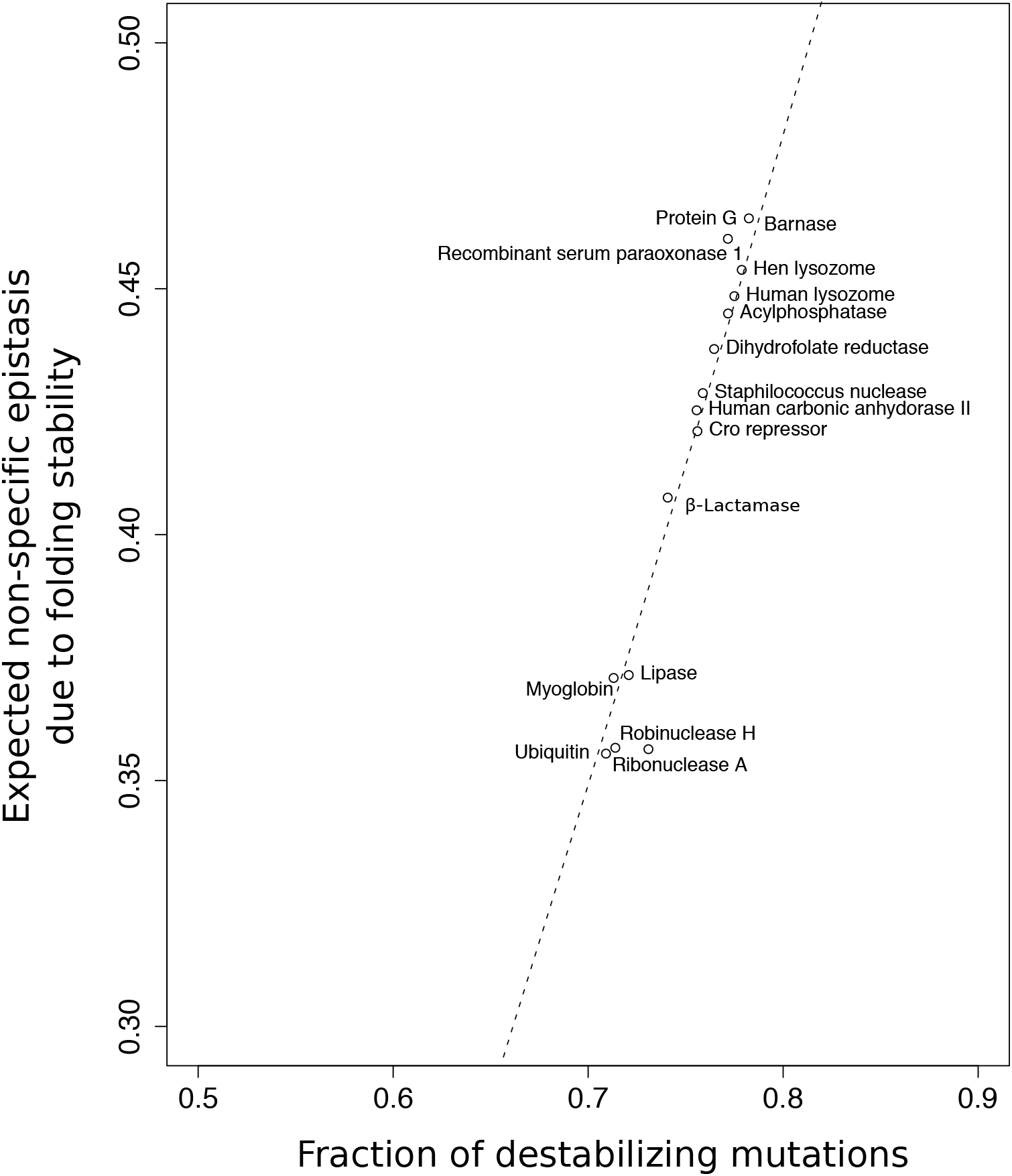
Expected epistasis imposed by folding stability increases by the fraction of destabilizing mutations. Dashed curve shows expected epistasis by folding stability when distribution of mutational effects on protein folding stability shifts towards destabilizing mutations. Expected epistasis for different globular proteins (filled circles) are shown. The line fitted to data has the equation: *epistasis*=1.52 × (*fraction of destabilizing mutations*) – 0.72. Fraction of destabilizing mutations is based on reference (<u>Tokuriki *et al*. 2007</u>).

### Theoretical analysis of non-specific epistasis to examine its dependence on fraction of destabilizing random mutations in a protein fold

The effect of random mutations on folding stability ΔΔ*G* affects evolutionary rates (Equation 4), and thus can influence the estimation of epistasis. Although the shape of the distribution of ΔΔ*G* due to random mutations is consistent across several types and diverse folds of proteins (<u>Tokuriki *et al*. 2007</u>), there are notable differences, such as the percentage of mutations that are stabilizing or destabilizing. To explore the robustness of estimates of epistasis on the distribution of arising random ΔΔ*G* in different protein folds, we resort to theoretical analysis of Equation 1. Specifically, rate of substitution is the product of the rate of mutation and the probability of fixation. Thus, for the non-synonymous rate d*N*=(µ*N*)Pfix, where µ is the mutation rate per generation. Since there are *N* birth events in a generation, *Nµ* is the number of random mutations per generation. For the synonymous mutation rate d*S*=(*µN*)(1/*N*), where the factor (1/N) is the probability of fixation for neutral mutation and in our model, synonymous mutations are neutral. Thus,

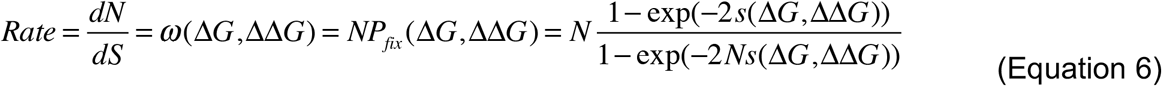

where we use **Equations 3** and **4** for the specific case of selection for folding stability. Equation 6 is the *dN*/*dS* for a random mutation that changes the wildtype folding stability, Δ*G*, by an amount ΔΔ*G*. The rate R_dN/dS_ is calculated multiple pairs of sequences in an MSA (Figure 1), thus it reflects the average *dN/dS* over many backgrounds and multiple arising mutations. Thus, we can arrive at an theoretical estimate of R_dN/dS_ by integrating over the distribution of wildtype folding stability Δ*G* and the distribution effects on folding stability due to random mutations ΔΔ*G*. The distribution P(ΔΔ*G*) is the probability distribution of mutational effects on folding stability known from large-scale mutational studies (<u>Alber 1989</u>; <u>Guerois *et al*. 2002</u>; <u>Tokuriki *et al*. 2007</u>; <u>Soskine and Tawfik 2010</u>), and has been parameterized for proteins of different folds. The distribution of background folding stability P(Δ*G*) is a consequence of mutation-selection balance on the protein folding fitness landscape (<u>Taverna and Goldstein 2002b</u>; <u>Taverna and Goldstein 2002a</u>; <u>Bloom *et al*. 2007</u>; <u>Zeldovich *et al*. 2007</u>; <u>Serohijos and Shakhnovich 2014b</u>). This distribution has also been documented experimentally from ~4000 proteins (<u>Bava *et al*. 2004</u>). For self-consistency, we numerically derive the distribution of folding stability stability p(Δ*G*) under mutation-selection balance using the parameters used in our sequence simulation (see Methods). Altogether, because the distributions P(ΔΔ*G*) and P(Δ*G*) are well-determined, we can arrive at an estimate of *R*_d*N*/d*S*_.

Next, we seek a theoretical estimate of the mutational usage, *R_u_*. To do so, we note that each protein sequence in an MSA corresponds to a Δ*G* value in the distribution P(Δ*G*). That is, each sequence in an MSA is a random sampling of the P(Δ*G*) distribution. Since *R_u_* assumes site-independence (<u>Breen *et al*. 2012</u>), we can consider each site to follow the distribution P(Δ*G*). Similarly, for a given site, each amino acid is a random sampling of the P(Δ*G*) distribution. In the language of molecular evolution, the P(Δ*G*) distribution may be considered as the site-equilibrium frequency in analogy to site-independence models of amino acids. Additionally, we note that in our simple model of selection for folding stability, in the regime of very high stability (Δ*G* < -20 kcal/mol), more mutations are allowed and amino acid usage (Figure S2, **black line**). Thus, curating an MSA of *k* orthologous sequences is sampling the P(Δ*G*) distribution *k* times (Figure S2). From the MSA, the value *u* is an upperbound estimate of mutational usage because it counts the number of *unique* amino acid in a site irrespective of frequency. The equivalent in our sampling method is the highest *u* from the *k* sampling of the P(Δ*G*) distribution (Figure S2). We used a cut-off of P(∆G)=0.001 and estimated *R_u_* as the evolutionary rate at highest stability at this probability:

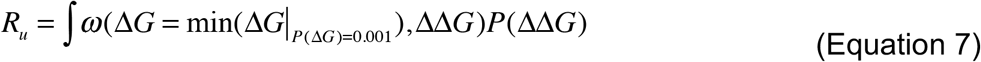

To check the consistency between simulations and our theoretical approach, we calculated *R_u_* for different number of sequences and also from Equation 6. Figure S3 shows the excellent agreement (r^2^=0.97, p-value < 10^−16^) between *R_u_* calculated from theory and simulations. The higher values of *R_u_* is calculated form theory is because of conceivable higher d*N* within the theoretical approach. Since all the calculations in the theoretical model are done with a continuous ∆∆G distribution, this condition is only achieved when all residues or at least a viable fraction have been mutated once in the sequence-explicit approach. Using both *R_dN_*_/*dS*_ and *R_u_* calculated from theory, we arrived at ε ~ 0.35−0.45, which is consistent with our results from explicit sequence simulation (Figure 2).

The theoretical approach enables us to investigate the sensitivity of estimated epistasis to the distribution of mutational effects (Table 1 and **Figure 5**), in particular, the fraction of mutations that are destabilizing. The values for averages and standard deviations of *P*(ΔΔ*G*) distributions are reported values for real proteins from large-scale mutagenesis studies (<u>Tokuriki *et al*. 2007</u>). We plotted the expected percentage of epistasis imposed by selection for folding stability of different globular proteins in **Figure 5** (see Table S3 for details). For example, average and standard deviation of P(ΔΔG>0) (bi-gaussian distribution (see Methods)) can vary from (1.33, 0.42) ± (1.64, 0.83) in Ubiquitin to (3.02,2.29) ± (0.76, 1.12) in Human lysozyme giving rise to ε=0.35 and ε=0.46 for the two proteins within our approach, respectively. As shown in Table 1, minor changes in P(ΔΔG>0) from 0.62 to 0.72 changes percentage of epistasis from 12% to 32%. In a hypothetical protein when the fraction of stabilizing and destabilizing mutations are almost equal ε is negligible.

**Table 1.**
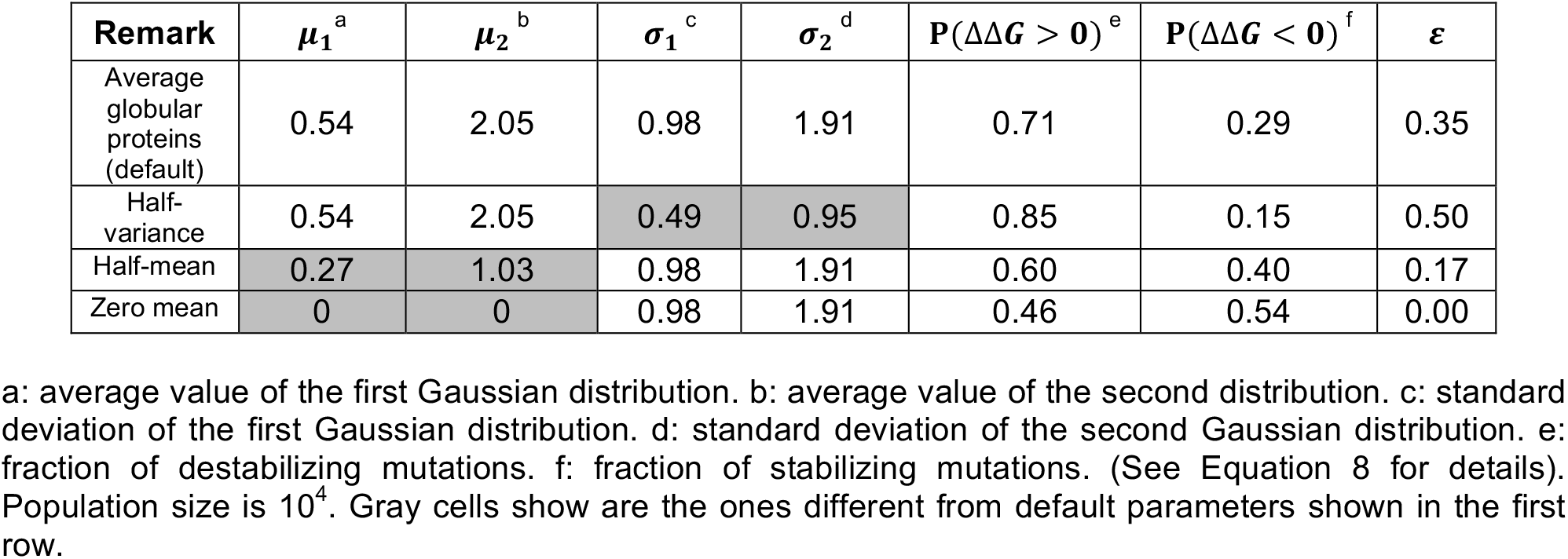
Sensitivity of estimated epistasis to parameters of distribution of mutational effects on protein folding stability.

## DISCUSSION

To what extent does the estimated ~30% epistasis agree with estimates from proteome-wide observations? Although several multiple factors that potentially contributing to epistasis, some of them beyond the property of one single gene (protein-protein interaction, centrality in a metabolic pathway, or genetic interactions), selection for folding stability have a major role, as quantified in this work. Since, folding stability is a primary selective force in protein evolution, and our model is based on a simple two-state folding thermodynamics, our estimate of ~30% epistasis sets a lower limit for epistasis experienced by real proteins. The higher limit for epistasis in molecular evolution might be ~90% epistasis as reported by Breen et al. (<u>Breen *et al*. 2012</u>).

Additionally, estimating epistasis in long-term protein evolution by comparing *R_dN/dS_* and *R_u_* from a multiple sequence alignment is simple, but not without potential complications. Plotkin and co-workers (<u>McCandlish *et al*. 2013</u>) argued that correcting the approach of Breen et al. (<u>Breen *et al*. 2012</u>) by using a distribution of selection coefficients for nonsynonymous substitutions could yield to a lower estimate of epistasis. In our approach, each nonsynonymous substitution would indeed have a different selection coefficient which not only depends on the effect size of mutation but also on the background stability in which it occurs. Therefore, the impact of nonsynonymous mutations is modeled more realistically by our approach, and the 30% estimate from folding stability is robust.

Furthermore, in line with two recent experimental systematic study of epistasis in protein G domain 1 (GB1) (<u>Olson *et al*. 2014</u>) and Hsp90 in yeast (<u>Bank *et al*. 2015</u>), we show that negative epistasis is the major type of epistasis under selection for PFS. Negative epistasis is mainly caused by sampling curved parts of fitness landscape, i.e., lower stabilities in the folding stability landscape. As illustrated in Figure S4, any factor that increases further sampling of the curvature of fitness landscape would increase epistasis. Indeed, adding other biophysical properties that could be relevant to fitness such as activity, dynamics, and binding to other proteins will only increase the ruggedness of the landscape, and hence the estimated epistasis.

## METHODS

### Protein evolution model and simulated phylogenetic tree

Protein sequences were evolved using a Wright-Fisher sampling approach with two fitness functions described by equations (4) and (5). In brief, codons were randomly mutated in one of the sites and once a nonsynonymous substitution arose, the relevant probability of fixation was calculated by the following equation:

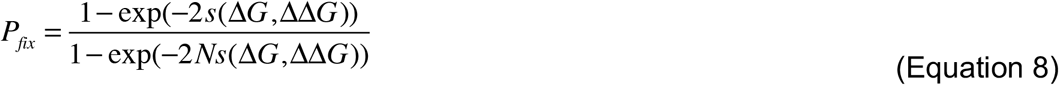

while *s*(∆*G*, ∆∆*G*) depends on the choice of fitness function. We assumed an effective monoclonal population size of N=10^4^. Initial sequence was taken to be that of *dihydrofolate reductase*, DHFR, taken from *Candida Albicans* with PDB ID=1AI9_A_1 (<u>Whitlow *et al*. 1997</u>). As described previously, the effect of mutations on protein stability was calculated using ERIS force field. To simulate sequence divergence, we simulated 2000 trajectories all started from an ancestral sequence and sampled 100 time points every 10^5^ mutational attempts. The 2000 sequences at each sampling point were regarded as orthologues used for estimation of epistasis.

To do so, we performed sequence-explicit simulations on the protein folding landscape (Figure 1) (Methods). To do so, we simulated dihydrofolate reductase (DHFR) sequences evolved under selection for thermodynamic stability (see Methods). DHFR is an integral protein in DNA nucleotide synthesis, as it converts dihydrofolic acid to tetrahydrofolic acid (<u>Fierke *et al*. 1987</u>) and therefore has a highly conserved function across organisms (<u>Hecht *et al*. 2011</u>). Several studies have shown that selection for thermodynamic stability might be the primary driving force in the evolution of DHFR (<u>Bershtein *et al*. 2012</u>; <u>Bershtein *et al*. 2015a</u>).

### Evolutionary rate estimation and pair epistasis

The pairwise rate of evolution, *R*_dN/dS_ =d*N*/d*S* and d*S* of DHFR sequences were estimated by Maximum-likelihood (ML) codon based model in codeml within PAML suite (<u>Yang 2007</u>). We estimated codon frequencies from the products of the average observed nucleotide frequencies in three codon positions (F3X4).

### Estimating limiting distribution of protein stabilities

To estimate epistasis using our numerical approach, we used distribution of mutational effects on protein stability, P(ΔΔG), from the extensive study of thousands of mutants in different proteins (<u>Tokuriki *et al*. 2007</u>):

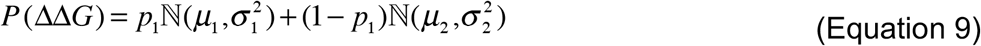

where *p*_1_ is the weights of first distribution, *μ*_1_, *μ*_2_, *σ*_1_ and *σ*_2_ are the average values and standard deviations of each Gaussian distribution found to be 0.53 ± 0.12, 0.56 ± 0.12, 1.96 ± 0.53, 0.90 ± 0.16 and, 1.93 ± 0.29 respectively. The distribution of background stabilities, P(ΔG), however, is a limiting distribution resulting from mutational supply and selection. We have previously used a numerical algorithm to obtain limiting distribution of protein stabilities under mutation-selection balance (<u>Kepp and Dasmeh 2014</u>). In brief, we start with an initial distribution *P*(Δ*G*)_*t* = 0_, which is simply the distribution of mutational effects centered on an initial stability, ΔG_int_. This distribution is then updated iteratively as:

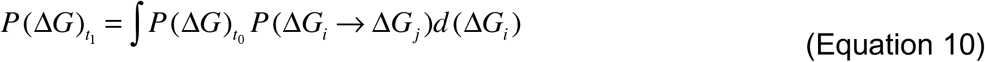

Here *P*(Δ*G*_i_ → Δ*G_j_*) is the transition probability of a protein from ∆*G_i_* to ∆*G_j_* which is the product of arising mutations and their fixation probability:

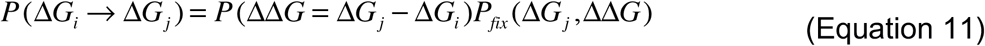

This approach is equivalent to locally weighted random walk sampling where each stability is sampled per the known distribution at time t_n_ giving rise to the distribution at time t_n+1_. Each time step in this numerical scheme is one mutational step. This procedure is continued iteratively while the convergence to a limiting distribution is reached judged by Kolmogorov-Smirinov two-sample test (Figure S5).

## Estimating the effect of point mutations on protein folding stability

To estimate the effect of site variations in mutational effects, i.e., ΔΔG=ΔG_mutated_ - ΔG_premutated_, we took ΔΔG values calculated for Dihydrofolate reductase (<u>Serohijos and Shakhnovich 2014a</u>). In brief, ΔΔGs are calculated using the flexible-back bone method of the ERIS algorithm (<u>Yin *et al*. 2007</u>). All side chains within 10Å of the mutated site were optimized and all dihedrals were relaxed to minimize backbone strain. This approach will give us a (Sequence length × 20 aa) matrix of ΔΔG values which is used for estimating the contribution of single sites to non-specific epistasis.

**Figure S1.**
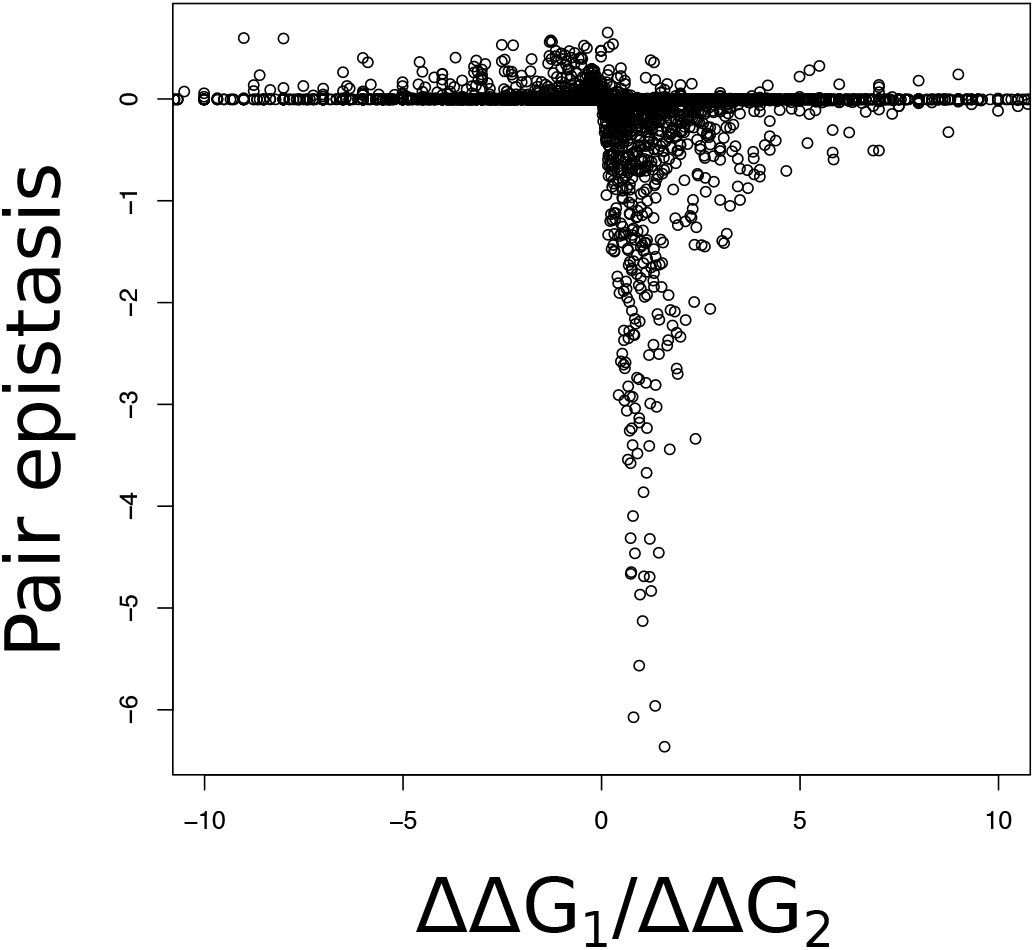
Pair epistasis (Equation 8 in the main text) versus the ratio of stability effect of two pair substitutions.

**Figure S2.**
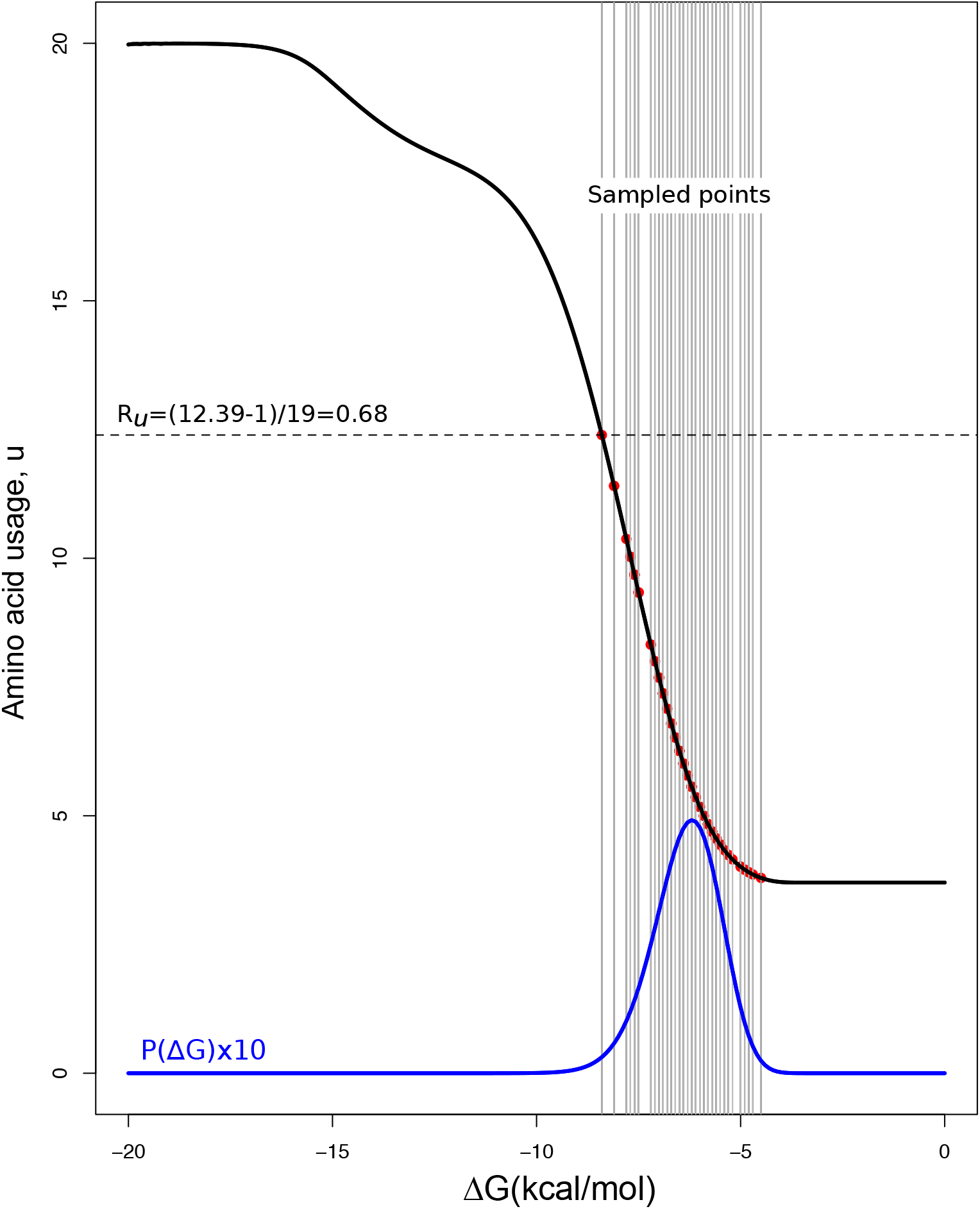
Fraction of fixed amino acids at different stabilities. Fixation is counted when N×P_fix_ ≥ 1. Red points represent 100 sampled stabilities and the vertical lines show the corresponding fraction of fixed amino acids. R_u_ is calculated as the fraction of fixed amino acids at the highest sampled stability. The blue plot shows 10×P(**∆**G). The factor 10 is multiplied by P(**∆**G) to scale the distribution for the presentation purpose.

**Figure S3.**
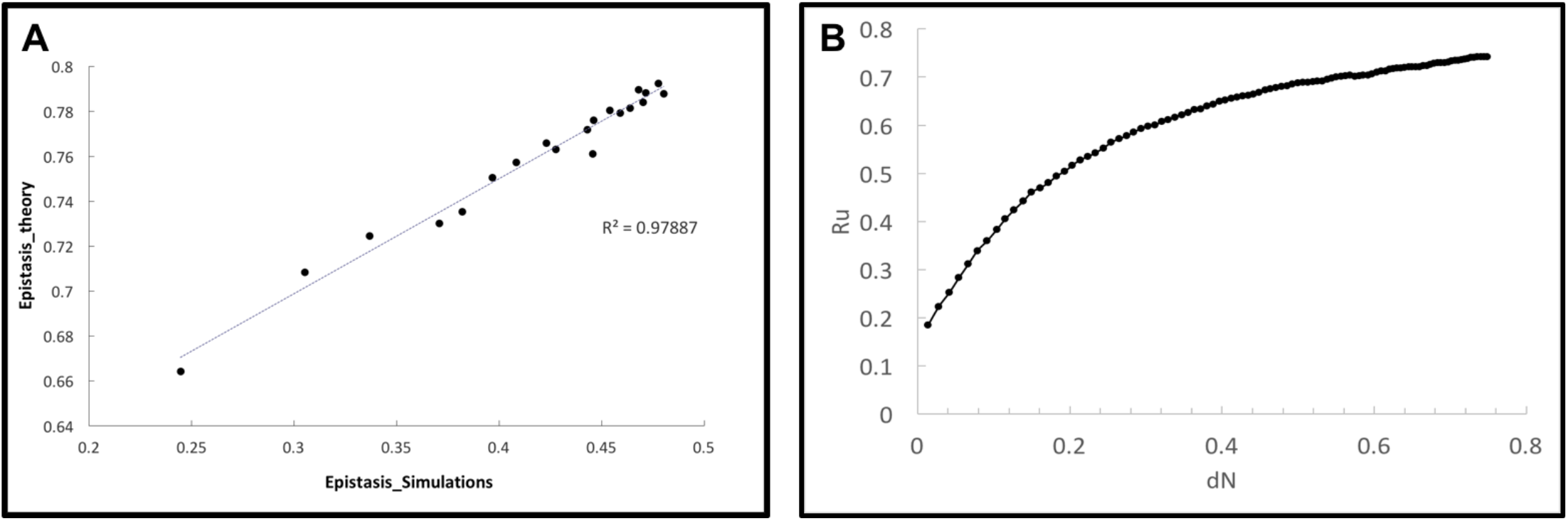
**(A)** Calculated epistasis by the theoretical approach versus simulation (R2=0.97 and p-value<10^−16^ using Wilcoxon signed-rank test). **(B)** R_u_ versus normalized nonsynonymous substitution rate by nonsynonymous sites, d*N*.

**Figure S4.**
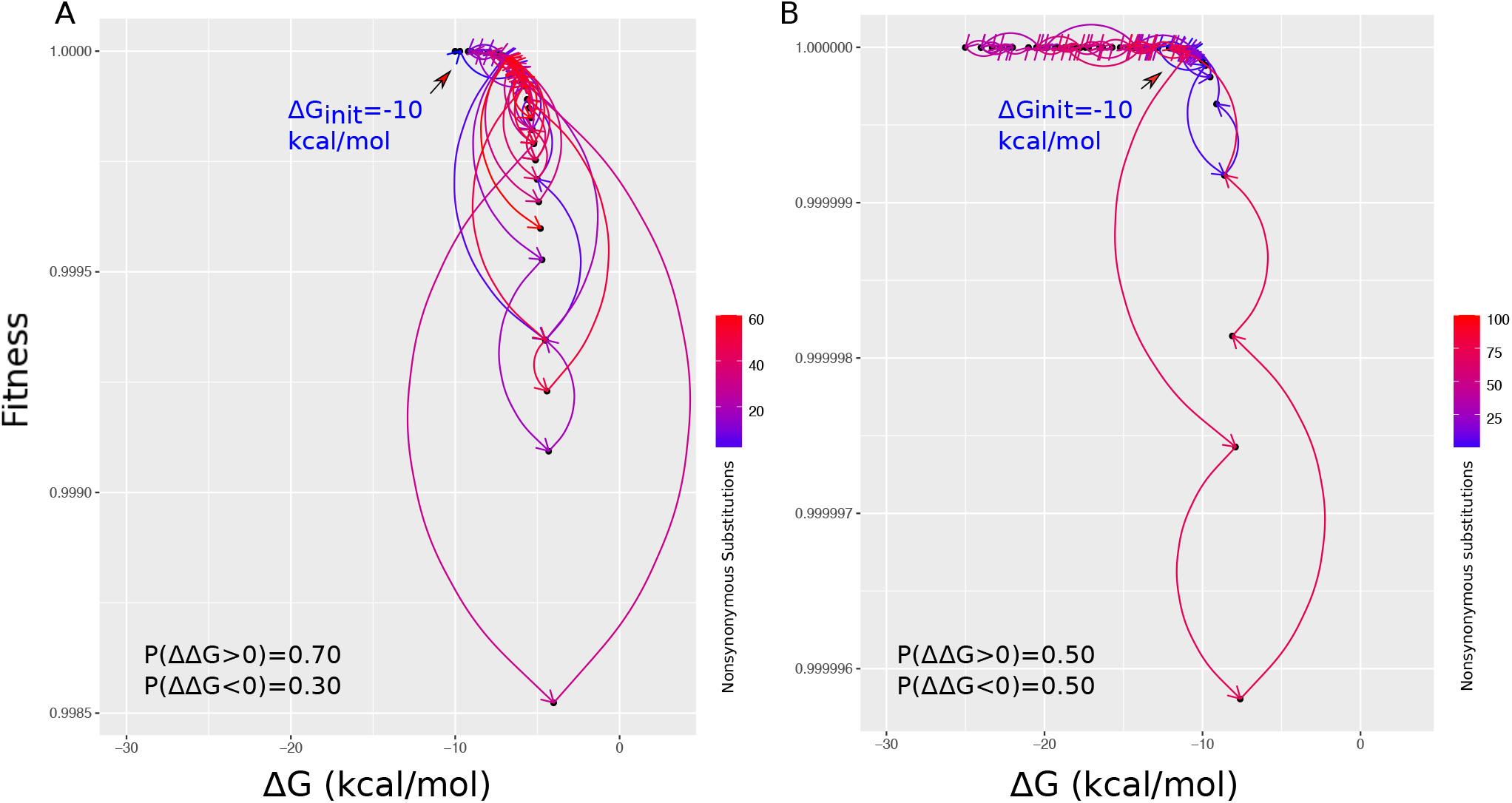
Destabilization shifts protein to more curved parts of fitness landscape and hence higher epistasis. **(A)** Evolutionary trajectory of an ancestral protein with stability of -10 kcal/mol when fraction of destabilizing and stabilizing mutations are 0.7 and 0.3 respectively. **(B)** the same plot in A with the difference in the fraction of destabilizing and stabilizing mutations to be equal. In both plots, orders of fixed mutations are depicted from blue to red. Fitness is proportional to F_nat_, number of folded copies of proteins per cell.

**Figure S5.**
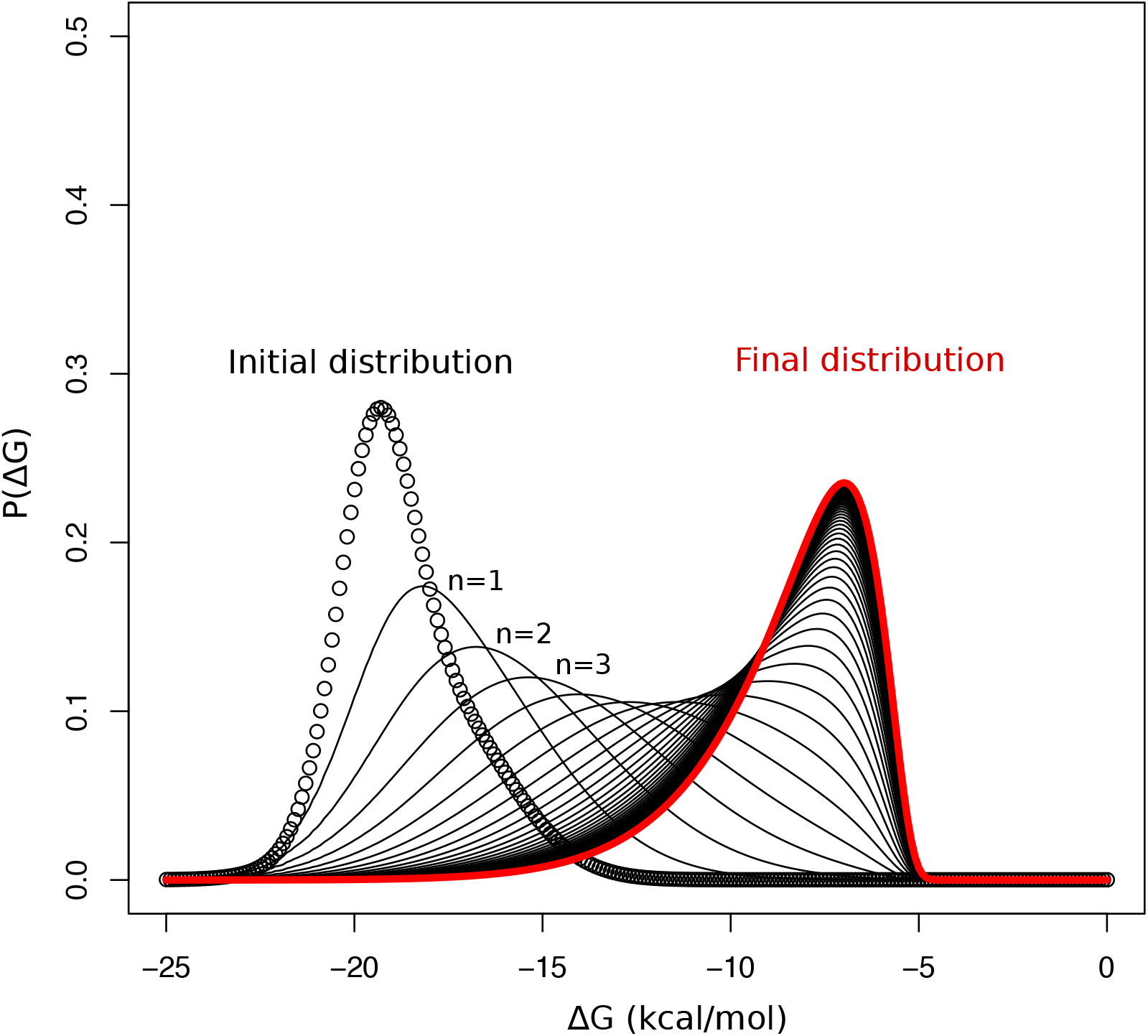
Evolution of distribution of protein stabilities within the numerical approach (equations 8 and 9 in the main text). The initial distribution is the distribution of mutational effects on a WT protein with ∆G = -20 kcal/mol without selection. The mutational steps are shown as n=1, n=2 and n=3 representing distribution of protein stabilities after one, two and three mutations respectively. The final distribution is shown in red which is the balance between mutation (i.e., causing destabilization) and selection for stabilizing mutations.

**Table S1.** Mutational usage, *u*, d*N*, d*S, R*_u_ and R_dN/dS_ for 2000 simulated trajectories sampled at 100 time points (each time point= 10N where N is the population size).

| Time point | $u$ | $dN$ | $dS$ | $R_u$ | $R_{dN/dS}$ |
| --- | --- | --- | --- | --- | --- |
| 1 | 4.4948 | 0.0134 | 0.0362 | 0.1839 | 0.3701 |
| 2 | 5.2396 | 0.0279 | 0.0720 | 0.2231 | 0.3871 |
| 3 | 5.7865 | 0.0415 | 0.1087 | 0.2519 | 0.3821 |
| 4 | 6.3802 | 0.0544 | 0.1447 | 0.2832 | 0.3761 |
| 5 | 6.9167 | 0.0667 | 0.1801 | 0.3114 | 0.3706 |
| 6 | 7.4427 | 0.0785 | 0.2163 | 0.3391 | 0.3629 |
| 7 | 7.8229 | 0.0910 | 0.2526 | 0.3591 | 0.3604 |
| 8 | 8.2760 | 0.1040 | 0.2890 | 0.3829 | 0.3598 |
| 9 | 8.6927 | 0.1152 | 0.3244 | 0.4049 | 0.3552 |
| 10 | 9.0469 | 0.1266 | 0.3614 | 0.4235 | 0.3502 |
| 11 | 9.3906 | 0.1385 | 0.4002 | 0.4416 | 0.3460 |
| 12 | 9.7500 | 0.1498 | 0.4362 | 0.4605 | 0.3434 |
| 13 | 9.9271 | 0.1613 | 0.4733 | 0.4698 | 0.3409 |
| 14 | 10.1250 | 0.1721 | 0.5108 | 0.4803 | 0.3369 |
| 15 | 10.3854 | 0.1824 | 0.5487 | 0.4940 | 0.3324 |
| 16 | 10.5729 | 0.1932 | 0.5866 | 0.5038 | 0.3293 |
| 17 | 10.8177 | 0.2034 | 0.6240 | 0.5167 | 0.3260 |
| 18 | 11.0260 | 0.2136 | 0.6599 | 0.5277 | 0.3237 |
| 19 | 11.1510 | 0.2235 | 0.7013 | 0.5343 | 0.3186 |
| 20 | 11.3073 | 0.2338 | 0.7405 | 0.5425 | 0.3158 |
| 21 | 11.4948 | 0.2440 | 0.7802 | 0.5524 | 0.3128 |
| 22 | 11.7135 | 0.2542 | 0.8185 | 0.5639 | 0.3106 |
| 23 | 11.8646 | 0.2647 | 0.8563 | 0.5718 | 0.3091 |
| 24 | 11.9740 | 0.2749 | 0.8960 | 0.5776 | 0.3068 |
| 25 | 12.1250 | 0.2843 | 0.9401 | 0.5855 | 0.3024 |
| 26 | 12.2656 | 0.2936 | 0.9784 | 0.5929 | 0.3000 |
| 27 | 12.3646 | 0.3027 | 1.0224 | 0.5981 | 0.2960 |
| 28 | 12.4010 | 0.3118 | 1.0686 | 0.6001 | 0.2917 |
| 29 | 12.5365 | 0.3207 | 1.1159 | 0.6072 | 0.2874 |
| 30 | 12.6146 | 0.3295 | 1.1715 | 0.6113 | 0.2812 |
| 31 | 12.7083 | 0.3381 | 1.2283 | 0.6162 | 0.2753 |
| 32 | 12.7969 | 0.3473 | 1.2925 | 0.6209 | 0.2687 |
| 33 | 12.8906 | 0.3552 | 1.3587 | 0.6258 | 0.2614 |
| 34 | 13.0052 | 0.3637 | 1.4270 | 0.6319 | 0.2549 |
| 35 | 13.0365 | 0.3720 | 1.5076 | 0.6335 | 0.2467 |
| 36 | 13.1510 | 0.3802 | 1.5994 | 0.6395 | 0.2377 |
| 37 | 13.2292 | 0.3885 | 1.6938 | 0.6436 | 0.2293 |
| 38 | 13.3385 | 0.3966 | 1.8081 | 0.6494 | 0.2193 |
| 39 | 13.3958 | 0.4045 | 1.9316 | 0.6524 | 0.2094 |
| 40 | 13.4531 | 0.4124 | 2.0540 | 0.6554 | 0.2008 |
| 41 | 13.4948 | 0.4200 | 2.2002 | 0.6576 | 0.1909 |
| 42 | 13.5417 | 0.4276 | 2.3312 | 0.6601 | 0.1834 |
| 43 | 13.5833 | 0.4348 | 2.4917 | 0.6623 | 0.1745 |
| 44 | 13.6250 | 0.4416 | 2.6533 | 0.6645 | 0.1664 |
| 45 | 13.6979 | 0.4490 | 2.8194 | 0.6683 | 0.1593 |
| 46 | 13.7813 | 0.4566 | 3.0083 | 0.6727 | 0.1518 |
| 47 | 13.8229 | 0.4636 | 3.1939 | 0.6749 | 0.1452 |
| 48 | 13.8802 | 0.4707 | 3.3855 | 0.6779 | 0.1390 |
| 49 | 13.9271 | 0.4779 | 3.5768 | 0.6804 | 0.1336 |
| 50 | 13.9427 | 0.4856 | 3.8040 | 0.6812 | 0.1277 |
| 51 | 14.0260 | 0.4922 | 3.9935 | 0.6856 | 0.1233 |
| 52 | 14.0573 | 0.4994 | 4.1985 | 0.6872 | 0.1190 |
| 53 | 14.0833 | 0.5063 | 4.3951 | 0.6886 | 0.1152 |
| 54 | 14.0990 | 0.5131 | 4.6142 | 0.6894 | 0.1112 |
| 55 | 14.1146 | 0.5192 | 4.8394 | 0.6902 | 0.1073 |
| 56 | 14.1406 | 0.5253 | 5.0564 | 0.6916 | 0.1039 |
| 57 | 14.1458 | 0.5319 | 5.2396 | 0.6919 | 0.1015 |
| 58 | 14.1979 | 0.5382 | 5.4282 | 0.6946 | 0.0992 |
| 59 | 14.2552 | 0.5441 | 5.6198 | 0.6976 | 0.0968 |
| 60 | 14.2917 | 0.5501 | 5.7863 | 0.6996 | 0.0951 |
| 61 | 14.3177 | 0.5565 | 5.9731 | 0.7009 | 0.0932 |
| 62 | 14.3490 | 0.5621 | 6.1320 | 0.7026 | 0.0917 |
| 63 | 14.3698 | 0.5683 | 6.3050 | 0.7037 | 0.0901 |
| 64 | 14.3281 | 0.5746 | 6.5007 | 0.7015 | 0.0884 |
| 65 | 14.3385 | 0.5801 | 6.6560 | 0.7020 | 0.0872 |
| 66 | 14.3698 | 0.5856 | 6.8187 | 0.7037 | 0.0859 |
| 67 | 14.3802 | 0.5921 | 6.9661 | 0.7042 | 0.0850 |
| 68 | 14.4271 | 0.5972 | 7.1060 | 0.7067 | 0.0840 |
| 69 | 14.4792 | 0.6034 | 7.2342 | 0.7094 | 0.0834 |
| 70 | 14.5260 | 0.6090 | 7.3925 | 0.7119 | 0.0824 |
| 71 | 14.5365 | 0.6149 | 7.5238 | 0.7124 | 0.0817 |
| 72 | 14.5990 | 0.6204 | 7.6460 | 0.7157 | 0.0811 |
| 73 | 14.6354 | 0.6249 | 7.7511 | 0.7177 | 0.0806 |
| 74 | 14.6615 | 0.6300 | 7.8720 | 0.7190 | 0.0800 |
| 75 | 14.6458 | 0.6350 | 7.9788 | 0.7182 | 0.0796 |
| 76 | 14.6667 | 0.6402 | 8.0832 | 0.7193 | 0.0792 |
| 77 | 14.6927 | 0.6456 | 8.1622 | 0.7207 | 0.0791 |
| 78 | 14.6875 | 0.6501 | 8.2432 | 0.7204 | 0.0789 |
| 79 | 14.6979 | 0.6548 | 8.3070 | 0.7209 | 0.0788 |
| 80 | 14.6979 | 0.6596 | 8.3776 | 0.7209 | 0.0787 |
| 81 | 14.7344 | 0.6649 | 8.4354 | 0.7229 | 0.0788 |
| 82 | 14.7552 | 0.6691 | 8.5072 | 0.7240 | 0.0787 |
| 83 | 14.7969 | 0.6737 | 8.5862 | 0.7262 | 0.0785 |
| 84 | 14.8385 | 0.6783 | 8.6573 | 0.7283 | 0.0784 |
| 85 | 14.8646 | 0.6832 | 8.7226 | 0.7297 | 0.0783 |
| 86 | 14.8594 | 0.6879 | 8.7751 | 0.7294 | 0.0784 |
| 87 | 14.8698 | 0.6924 | 8.8326 | 0.7300 | 0.0784 |
| 88 | 14.8958 | 0.6973 | 8.8836 | 0.7314 | 0.0785 |
| 89 | 14.9219 | 0.7016 | 8.9306 | 0.7327 | 0.0786 |
| 90 | 14.9479 | 0.7063 | 8.9804 | 0.7341 | 0.0787 |
| 91 | 14.9583 | 0.7105 | 9.0179 | 0.7346 | 0.0788 |
| 92 | 14.9740 | 0.7146 | 9.0494 | 0.7355 | 0.0790 |
| 93 | 14.9948 | 0.7191 | 9.0898 | 0.7366 | 0.0791 |
| 94 | 15.0156 | 0.7233 | 9.1252 | 0.7377 | 0.0793 |
| 95 | 15.0625 | 0.7273 | 9.1542 | 0.7401 | 0.0794 |
| 96 | 15.0833 | 0.7316 | 9.1820 | 0.7412 | 0.0797 |
| 97 | 15.0885 | 0.7358 | 9.2186 | 0.7415 | 0.0798 |
| 98 | 15.0885 | 0.7403 | 9.2497 | 0.7415 | 0.0800 |
| 99 | 15.0938 | 0.7445 | 9.2742 | 0.7418 | 0.0803 |
| 100 | 15.0885 | 0.7483 | 9.3081 | 0.7415 | 0.0804 |

**Table S2.** Mutational usage, *u*, d*N*, d*S, R*_u_ and R_dN/dS_ for different number of sampled sequences in simulations and theory.

| Number of sequences | $u$ | $dN$ | $dS$ | $dNdS_{simul}$ | $dNdS_{theory}$ | $R_u_{simul}$ | $R_u_{theory}$ |
| --- | --- | --- | --- | --- | --- | --- | --- |
| 100 | 5.6458 | 0.1738 | 0.4993 | 0.3482 | 0.4256 | 0.2445 | 0.6643 |
| 200 | 6.7969 | 0.1742 | 0.5103 | 0.3413 | 0.4252 | 0.3051 | 0.7085 |
| 300 | 7.3958 | 0.1720 | 0.5105 | 0.3369 | 0.4227 | 0.3366 | 0.7247 |
| 400 | 8.0417 | 0.1727 | 0.5090 | 0.3394 | 0.4229 | 0.3706 | 0.7303 |
| 500 | 8.2552 | 0.1720 | 0.5104 | 0.3370 | 0.4239 | 0.3819 | 0.7354 |
| 600 | 8.5365 | 0.1723 | 0.5122 | 0.3365 | 0.4238 | 0.3967 | 0.7505 |
| 700 | 8.7552 | 0.1722 | 0.5086 | 0.3385 | 0.4247 | 0.4082 | 0.7575 |
| 800 | 9.0365 | 0.1722 | 0.5058 | 0.3405 | 0.4240 | 0.4230 | 0.7660 |
| 900 | 9.1250 | 0.1727 | 0.5089 | 0.3394 | 0.4230 | 0.4276 | 0.7632 |
| 1000 | 9.4167 | 0.1724 | 0.5108 | 0.3375 | 0.4240 | 0.4430 | 0.7719 |
| 1100 | 9.4635 | 0.1729 | 0.5084 | 0.3401 | 0.4240 | 0.4454 | 0.7613 |
| 1200 | 9.4740 | 0.1719 | 0.5122 | 0.3356 | 0.4243 | 0.4460 | 0.7762 |
| 1300 | 9.6250 | 0.1726 | 0.5123 | 0.3369 | 0.4241 | 0.4539 | 0.7806 |
| 1400 | 9.7188 | 0.1728 | 0.5098 | 0.3390 | 0.4237 | 0.4589 | 0.7793 |
| 1500 | 9.8125 | 0.1716 | 0.5098 | 0.3366 | 0.4227 | 0.4638 | 0.7815 |
| 1600 | 9.8906 | 0.1714 | 0.5113 | 0.3352 | 0.4245 | 0.4679 | 0.7898 |
| 1700 | 9.9323 | 0.1719 | 0.5107 | 0.3366 | 0.4239 | 0.4701 | 0.7842 |
| 1800 | 9.9583 | 0.1720 | 0.5110 | 0.3366 | 0.4233 | 0.4715 | 0.7883 |
| 1900 | 10.0729 | 0.1723 | 0.5105 | 0.3375 | 0.4236 | 0.4775 | 0.7925 |
| 2000 | 10.1250 | 0.1721 | 0.5108 | 0.3369 | 0.4235 | 0.4803 | 0.7880 |

**Table S3.** Parameters of distribution of mutational effects for different globular proteins.

| Protein | $\mu_1^a$ | $\mu_2^b$ | $\sigma_1^c$ | $\sigma_2^d$ | $P_1^e$ | $P(\Delta\Delta G > 0)$ | % epistasis |
| --- | --- | --- | --- | --- | --- | --- | --- |
| Recombinant serum<br>paraoxonase 1 | 0.51 | 0.91 | 1.93 | 1.82 | 0.42 | 0.77 | 0.33 |
| Lipase | 0.57 | 1.18 | 2.27 | 2.05 | 0.67 | 0.72 | 0.25 |
| β-Lactamase | 0.58 | 1.11 | 2.36 | 1.84 | 0.67 | 0.74 | 0.29 |
| Human carbonic anhydrase II | 0.5 | 0.85 | 1.8 | 1.99 | 0.39 | 0.76 | 0.29 |
| Dihydrofolate reductase | 0.54 | 0.73 | 1.53 | 1.78 | 0.39 | 0.76 | 0.29 |
| Robonuclease H | 0.49 | 0.98 | 2.12 | 1.98 | 0.69 | 0.71 | 0.24 |
| Myoglobin | 0.31 | 0.84 | 1.53 | 1.57 | 0.48 | 0.71 | 0.25 |
| Staphilococcus nuclease | 0.58 | 0.68 | 1.3 | 1.71 | 0.3 | 0.76 | 0.27 |
| Human lysosome | 0.76 | 1.12 | 3.02 | 2.29 | 0.7 | 0.77 | 0.34 |
| Hen lysosome | 0.81 | 1.16 | 3 | 2.43 | 0.68 | 0.78 | 0.34 |
| Ribonuclease A | 0.59 | 0.83 | 1.76 | 2.56 | 0.55 | 0.73 | 0.21 |
| Barnase | 0.59 | 0.84 | 2.08 | 1.93 | 0.51 | 0.78 | 0.34 |
| Acylphosphatase | 0.56 | 0.8 | 1.93 | 1.83 | 0.56 | 0.77 | 0.32 |
| Ubiquitin | 0.42 | 0.83 | 1.33 | 1.64 | 0.52 | 0.71 | 0.22 |
| Protein G | 0.59 | 0.84 | 2.08 | 1.93 | 0.51 | 0.78 | 0.34 |
| Cro repressor | 0.55 | 0.74 | 1.33 | 1.58 | 0.48 | 0.76 | 0.27 |

a: average value of the first Gaussian distribution. b: average value of the second distribution. c: standard deviation of the first Gaussian distribution. d: standard deviation of the second Gaussian distribution. e: fraction of destabilizing mutations. f: fraction of stabilizing mutations. (See Equation 8 for details). Population size is $10^4$ . Gray cells show are the ones different from default parameters shown in the first row. e: weight of first Gaussian distribution (see Equation 7 in the main text). Data is taken from (TOKURIKI et al. 2007).

